# IncC plasmid genome rearrangements influence the vertical and horizontal transmission tradeoff in *Escherichia coli*

**DOI:** 10.1101/2024.04.17.589899

**Authors:** Margaux Allain, Thibaut Morel-Journel, Bénédicte Condamine, Benoist Gibeaux, Benoit Gachet, Rémi Gschwind, Erick Denamur, Luce Landraud

## Abstract

It has been shown that an evolutionary tradeoff between vertical (host growth rate) and horizontal (plasmid conjugation) transmissions contribute to global plasmid fitness. As conjugative IncC plasmids are important for the spread of multidrug resistance (MDR), in a broad range of bacterial hosts, we investigated vertical and horizontal transmissions of two multidrug-resistant IncC plasmids according to their backbones and MDR-region rearrangements, upon plasmid entry into a new host. We observed plasmid genome deletions after conjugation in three diverse natural *Escherichia coli* clinical strains, varying from null to high number depending on the plasmid, all occurring in the MDR-region. The plasmid burden on bacterial fitness depended more on the strain background than on the structure of the MDR-region, deletions appearing to have no impact. Besides, we observed an increase in plasmid transfer rate, from ancestral host to new clinical recipient strains, when the IncC plasmid was rearranged. Finally, using a second set of conjugation experiments, we investigated the evolutionary tradeoff of the IncC plasmid during the critical period of plasmid establishment in *E. coli* K-12, by correlating the transfer rates of deleted or non-deleted IncC plasmids and their costs on the recipient strain. Plasmid deletions strongly improved conjugation efficiency with no negative growth effect. Our findings indicate that the flexibility of the MDR-region of the IncC plasmids can promote their dissemination, and provide diverse opportunities to capture new resistance genes. In a broader view, they suggest that the vertical-horizontal transmission tradeoff can be manipulated by the plasmid to improve its fitness.

## INTRODUCTION

IncC plasmids are large, low copy theta-replicating plasmids, and have an extremely broad-host range. First isolated in the late 1960s, they have been found among agricultural, veterinary, and human clinical isolates worldwide (1–3), including *Aeromonas ssp*, *Yersinia pestis*, *Photobacterium damselae subsp. piscicida*, *Klebsiella pneumoniae*, *Vibrio cholera*, *Escherichia coli*, and *Salmonella enterica* (4).

The term IncA/C has been frequently used to describe plasmids belonging to incompatibility groups A and C (2), linked to some entry-exclusion data which suggested that plasmids from these two groups were very closely related. Two distinct IncA/C lineages, designated A/C_1_ and A/C_2_, based on the differences in sequence identity of the plasmid amplicons, have been commonly used in the literature to distinguish the IncA plasmid from the IncC group (2, 5). However, recent works of Ambrose *et al.* demonstrated that this designation was incorrect, since IncA and IncC plasmids were in fact compatible (6). Using the term IncA/C complicates the description of the phylogeny and evolution of IncC plasmid backbone genes, between the oldest IncC plasmids and contemporary IncC plasmids. It is now well admitted that all IncC plasmids share a highly conserved backbone, and they can be broadly typed into two groups (namely, types 1 and 2), which diverged a long time ago, based on a few differences in their backbones (7). Most of the diversity in IncC plasmid genomes results from changes, confined to a few hotspots of integration, consisting of differences within horizontally acquired accessory genes (in particular the gain and deletion of resistance genes) and/or loss of adjacent backbone regions (3, 7–9). Some type 1/2-hybrid IncC plasmids have been described, arising from homologous recombination events between the two groups of IncC plasmids, or multireplicon composite IncC plasmids, resulting from the integration of a plasmid belonging to another incompatibility group into the IncC plasmid genome (9–11).

IncC conjugative plasmids are regarded as a considerable public health threat due to their ability to confer resistance to multiple antimicrobial agents and their contribution to the dissemination of resistance genes, such as genes encoding AmpC β-lactamases, extended-spectrum β-lactamases (ESBL) or carbapenemases, from long-term hosts to other reservoir bacteria (12–14). The evolution of multidrug resistance (MDR)-encoding modules, via loss and acquisition of diverse resistance cassettes within especially two large antibiotic resistance islands, at a specific location in the backbone (classically designated ARI-A and ARI-B regions), might contribute to the emergence of new antimicrobial resistance in new pathogens. Although the burden on host cells associated with large costly plasmid acquisition is expected to limit plasmid dissemination in the absence of positive selection, IncC plasmids are frequently described and stably maintained, in an extremely broad range of bacterial hosts. In a previous work, we observed that a costly IncC plasmid was stably maintained without undergoing modification in an *E. coli* laboratory adapted bacterium, while many modifications of the MDR-encoding module appeared immediately after conjugation (15). Two interdependent forces govern the success of multidrug-resistant plasmids: the vertical transmission through cell division and the horizontal transmission through conjugation. In fact, plasmids can exert a variable range of fitness cost on their hosts, resulting in plasmid-carrying cells being outcompeted by plasmid-free cells, with a subsequent impact on the vertical transmission of the plasmid in the absence of positive selection (16). Furthermore, plasmid conjugation constitutes a costly process which may slow host growth rates, in particular immediately after acquisition in a new host (17). Numerous observations in the literature have reported that the plasmid transfer system is often targeted in host-plasmid adaptation processes resulting in decreased conjugation (9, 18, 19). Plasmid costs and conjugation efficiencies depend on general plasmid properties, and an optimal balance between vertical and horizontal transmission modes has been observed to explain high plasmid fitness and prevalence of some plasmids (20).

To gain insights into the role of rearrangements in IncC plasmid fitness, we investigated vertical and horizontal transmission of these plasmids, according to their backbones and MDR-region rearrangements. More precisely, using an *in vitro* experimental approach and two different ESBL-encoding IncC plasmids (15), we determined plasmid rearrangement frequency, after conjugation experiments in three phylogenetically diverse clinical *E. coli* recipient strains from the Natural Isolates with Low Subcultures (NILS) collection (21). We then evaluated the impact of IncC plasmids on bacterial fitness, according to the backbone and MDR-region rearrangements, by determining the plasmid carriage burden on non-rearranged and rearranged plasmid-carrying transconjugants (TCs). We also evaluated plasmid stability in newly formed TCs according to the MDR-encoding structure, using an experimental evolution assay in the absence and presence of cefotaxime. Moreover, we compared conjugation frequencies to the clinical recipient strains, between plasmids with and without MDR-encoding module rearrangements. Lastly, we performed a second set of conjugation experiments, with *E. coli* K-12 as the recipient host, using the first generation of TCs as donor, to determine the global fitness of IncC plasmid according to the structure of the MDR-encoding region by correlating the plasmid acquisition cost for the K-12 recipient cell and the plasmid transfer rate to that cell (Figure S1).

## RESULTS

### Rational of the plasmids and strains used for conjugation

To understand the evolutionary mechanisms involved in the dissemination and stability of multidrug-resistant IncC plasmids in bacterial hosts, we used two plasmids (pRCS30 and pRCS46; for simplicity, we will refer to them as p30 and p46, respectively) from a previous collection of sequenced and well-characterized ESBL-encoding plasmids from human clinical *E. coli* strains (CB513 and CB195, respectively) (22). p30 carries a *bla*_CTX-M-14_ gene inserted between the *ter* and *kfr* genes, while p46 carries a *bla*_CTX-M-15_ gene, and the IS*Ec9*-*bla*_CTX-M-15_-ORF*477* transposition unit is inserted in the backbone of the plasmid, between a gene encoding a DNA methylase and a gene encoding a conserved protein of unknown function. Moreover, p46 has the particularity of being a multireplicon composite plasmid (IncC-IncR plasmid) that has integrated an IncR plasmid within the large MDR-encoding module, inserted at an integration spot interrupting the *rhs* gene. p46 and p30 belong to the type 2 lineage of IncC plasmids (Figure S2, S3). These two plasmids vary in size from 157 to 246 kb and carry multiple other resistance genes in addition to the *bla*_CTX-M_ genes (for additional information, see Materials and Methods and Table S1). We have observed that a small fraction of the ancestral CB195 donor strain (less than 3 % of the total bacterial population) harbored a plasmid, which has spontaneously lost a part of the MDR-encoding module after an *in vitro* experimental evolution (from 80 to 130 generations). Compared to p46, this plasmid (denoted as p46^r^) presents two deletions, a 13,345 bp fragment, just downstream of the *uvrD* gene, encoding the *aacC2-aph-sul2* cassette, which confers resistance to aminoglycosides, and a 48,745 bp fragment encoding the IncR backbone genes and their flanking regions (Table S1 and Figure S3).

We performed multiple independent transfers of these plasmids (p30 from CB513, and p46 and p46^r^ from CB195) in three clinical *E. coli* strains (NILS07, NILS70, and NILS71) made resistant to rifampicin, to obtain multiple independent TCs, denoted as TC-NILS. We selected clinical *E. coli* recipient strains for this study on the basis of their diverse phylogroups (A and B2), their sequence type representative of the current epidemiology, and their lack of antimicrobial resistance gene (for additional information, see Materials and Methods, Table S1).

### Analysis and frequency of plasmid rearrangements in clinical transconjugants

To identify plasmid mutations and rearrangements that may occur immediately after conjugation, we sequenced randomly selected TC-NILS obtained from independent conjugation assays. As no p30 rearrangement was observed in our previous work (15), we sequenced one TC-NILS obtained after conjugation of p30, and three to six TC-NILS after conjugation of p46, in each clinical recipient strain (n= 20, Table S2).

We used the open-source Breseq pipeline (23) and the ancestral p46 or p30 plasmids as reference genomes. In the first step, we excluded all single nucleotide polymorphisms (SNP), insertions or deletions detected comparing mapping reads of the plasmid references to the assembly of themselves. In the second step, we determined genetic differences occurring between the reference plasmids and the TC-NILS genomes.

For all TC-NILS harboring p30 (denoted as NILS-p30), we observed no plasmid DNA modification after conjugation. In contrast, for 12 of the 14 TC-NILS obtained from p46 plasmid conjugation, we detected deletions in the plasmid genome within a large 122,219 bp MDR-region inserted into the *rhs* gene (Figure 1 and Table S2). For 10 of these TC-NILS, the deletions occurred in two hotspots: an IS*26*-mediated deletion of a sequence encoding the complete backbone of the IncR plasmid and its flanking regions, and an IS*26*-mediated deletion of a sequence carrying aminoglycosides resistance module (*aacC2*-*aph-sul2* cassette) (Figure 1) (plasmids denoted as p46^r^, TC-NILS denoted as NILS-p46^r^). Four others TC-NILS from conjugation of the p46 plasmid harbored a plasmid without modification of the IncR plasmid genome and of the aminoglycosides resistance module (TC513 from NILS70 and TC549 from NILS71), or a plasmid without any deletions (TC569 and TC570, both from NILS07) (plasmids denoted as p46^nr^, TC-NILS denoted as NILS-p46^nr^). As expected, we observed a perfect correlation between the presence of antibiotic resistance genes and phenotypic antibiotic sensitivity results (Table S2).

**FIG 1:**
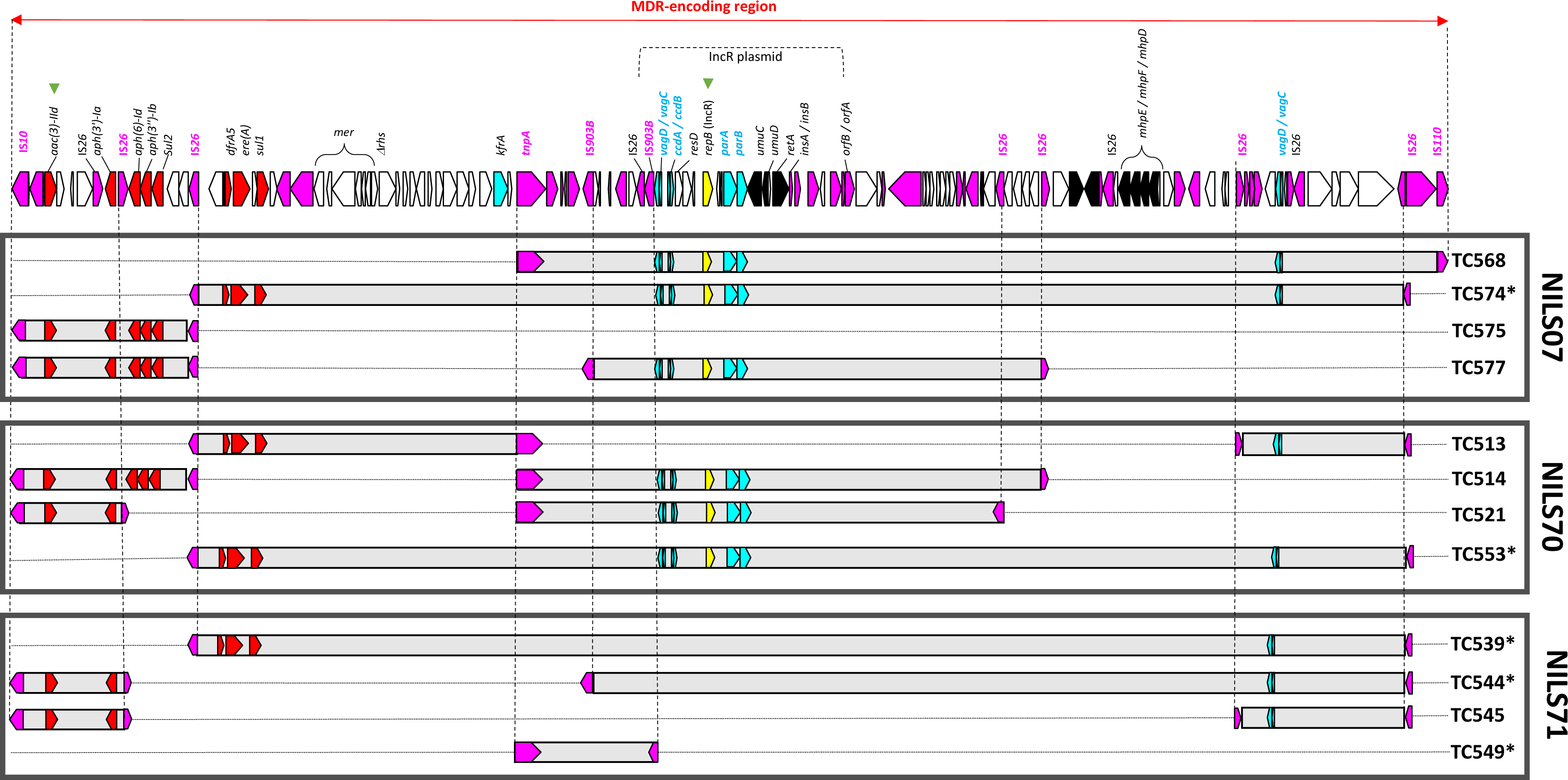
Representation of the MDR-encoding region of the p46 plasmid after conjugation of donor strain CB195 in 3 *E. coli* clinical isolates (NILS07, NILS70, and NILS71; indicated on the right). The linear map of the MDR-encoding region of the p46 plasmid, indicated by double arrow, is detailed and open reading frames are shown as arrows indicating the direction of transcription (yellow, replication; light blue, plasmid maintenance; red, resistance; black, metabolism; pink, mobile elements and white, hypothetical proteins). The IncR plasmid is indicated by dotted line. The regions involved in the rearrangements observed in each transconjugant (TC-NILS, referred on the right) are indicated by grey boxes and bordering IS are indicated by a dotted line. Resistance, maintenance and replication genes deleted in each transconjugant are represented in the grey boxes. Green triangles indicate the areas targeted in the rearrangement screening (see text). Asterisks indicate TC-NILS used in conjugation experiments in *E. coli* K-12 MG1655 (see text).

To test whether the high frequency of rearrangement events could be due to a hypermutator profile of the donor strains, we estimated mutation frequencies of strains CB513 (harboring p30) and CB195 (harboring p46) by monitoring their capacity to generate mutations conferring resistance to rifampicin or fosfomycin. We observed no significant differences compared with the non-hypermutator control strain (Figure S4). Moreover, we detected no mutation in the mismatch repair system in the genome of the strains, suggesting that they were non-hypermutator strains.

To estimate p46 plasmid rearrangement frequency after conjugation in clinical recipient strains, we screened a total of 104 TC-NILS from p46 plasmid conjugation (34, 38, and 32 TCs after conjugation in NILS07, NILS70, and NILS71, respectively), for phenotypic gentamicin resistance and for the presence of the IncR replicase-encoding gene (*repB* gene). Although our method could underestimate the p46 rearrangement frequency, we assumed that it allowed us to determine the minimum rate of plasmid genome modification, in accordance with our previous observations of conjugation-induced p46 deletions in these two hotspots. In agreement with the first data, we detected four profiles: gentamicin-resistant *repB*-positive strains (with no expected plasmid IncR backbone/aminoglycosides deletion), gentamicin-resistant *repB*-negative strains (with deletion of the IncR backbone), *repB*-positive strains with a restoration of gentamicin sensitivity (with deletion of the aminoglycosides cassette), and finally r*epB*-negative and gentamicin-sensitive strains (combining deletion of the IncR backbone and the aminoglycosides cassette). In NILS07, 5.9 % of the TCs had an IncR backbone deletion, 17.6 % of the TCs had their sensitivity to gentamicin restored and 2.9 % of the TCs had both. In NILS70, 10.5 % of the TCs had an IncR deletion, 7.9 % of the TCs had their sensitivity to gentamicin restored and 7.9 % of the TCs had both. In NILS71, 21.9 % of the TCs had an IncR deletion, 9.4 % of the TCs had their sensitivity to gentamicin restored and 6.3 % of the TCs tested had both.

Overall, the global observed deletion rates were high, between 26.3 % (NILS70) and 37.5 % (NILS71), and did not vary according to the recipient strain or the phylogroup (chi-squared test, *P* = 0.52) (Figure S5).

### Growth burden of IncC plasmids on *E. coli* clinical recipient strains

To evaluate the impact of IncC plasmid carriage on clinical *E. coli* recipient strains, and the growth burden of plasmid rearrangements on the clinical *E. coli* host, we measured maximum growth rates (MGRs) immediately after conjugation. These rates were analyzed using a multivariate regression, which provides quantitative estimates of the impact of specific factors in a single statistical model. According to the deletion profiles and the recipient strains, we tested 14 TC-NILS: 3 NILS-p30, 3 NILS-p46^nr^ without IncR backbone/aminoglycosides deletion, and 8 NILS-p46^r^ with IncR backbone/aminoglycosides deleted plasmid (Table S3). We compared the MGRs of the TC-NILS to the MGRs of the recipient plasmid-free strains (NILS07, NILS70, and NILS71) having been incubated concomitantly with the TC-NILS, and the MGRs of the TC-NILS with each other (Figure 2).

**FIG 2:**
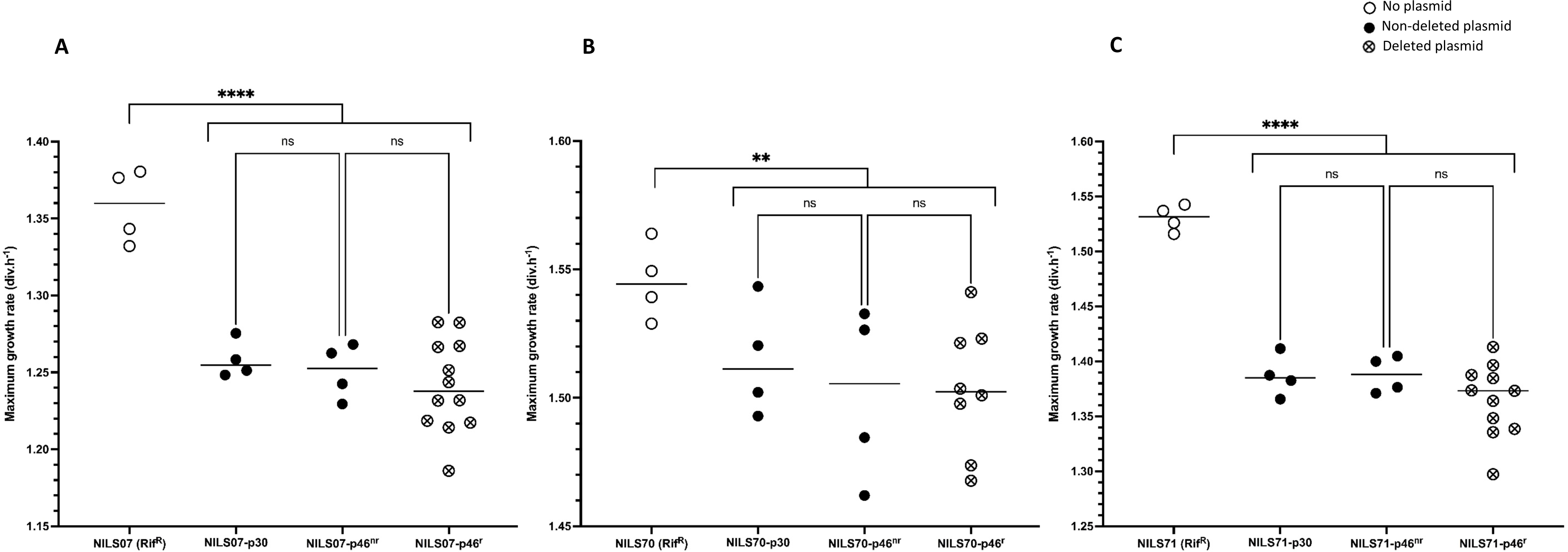
Scatter plot representation of the maximum growth rates (MGRs) of each NILS and TC-NILS. For each NILS, MGRs of NILS-Rif^R^ with no plasmid (empty circle), NILS-p30 (full circle), NILS-p46^nr^ (full circle), and NILS-p46^r^ (circle-cross) are represented. **(A)**, MGRs of plasmid-free and plasmid-carrying NILS07. **(B)**, MGRs of plasmid-free and plasmid-carrying NILS70. **(C)**, MGRs of plasmid-free and plasmid-carrying NILS71. Each dot represents the result of one experiment. Each central line represents the mean of all the experiments. All experiments were performed in three technical replicates. Asterisks indicate significant differences (**, *P* ≤ 0.01; ****, *P* ≤ 0.0001), ns = not significant.

The MGR of TC-NILS were significantly lower than those of the plasmid-free strains, but the difference varied with the strain considered. The greatest decrease was observed in TC-NILS71 (*β*_5_ = -0.15 div.h^-1^, *P* < 0.0001), followed by TC-NILS07 (*β*_3_ = -0.11 div.h^-1^, *P* <

0.0001) and TC-NILS70 (*β*_4_ = -0.04 div.h^-1^, *P* = 0.0098). No significant differences were observed between the MGRs of NILS-p30 and NILS-p46^nr^, or between the MGRs of NILS-p46^nr^ and NILS-p46^r^ (Figure 2, Table S3 and S4).

In conclusion, IncC plasmids systematically entail a cost to bacterial fitness, which varies more depending on the strain background than on MDR-encoding module structure, where deletions do not appear to have any impact on growth burden in our model.

### Effect of MDR-encoding region rearrangements on conjugation transfer frequency

To evaluate the impact of rearrangements on horizontal transmission of the p46 plasmid, we used CB195 cells harboring the plasmid which had spontaneously lost part of the MDR-encoding region (p46^r^) as donor, because the rearrangements were similar to those observed in TC-NILS after conjugation (deletion of the *aacC2-aph-sul2* cassette and the IncR region) (Figure S3), and we compared the transfer rate of p46 and p46^r^ from the CB195 ancestral strain to the three NILS clinical strains.

In NILS07, the transfer rate of the p46^r^ plasmid was 10.7 higher than that of the p46 plasmid (*P* = 0.047). In NILS70, the transfer rate of the p46^r^ plasmid was 3.2 higher than that of the p46 plasmid (*P* = 0.023). In NILS71, the transfer rate of the p46^r^ plasmid was 3.4 higher than that of the p46 plasmid (*P* = 0.028) (Figure 3).

**FIG 3:**
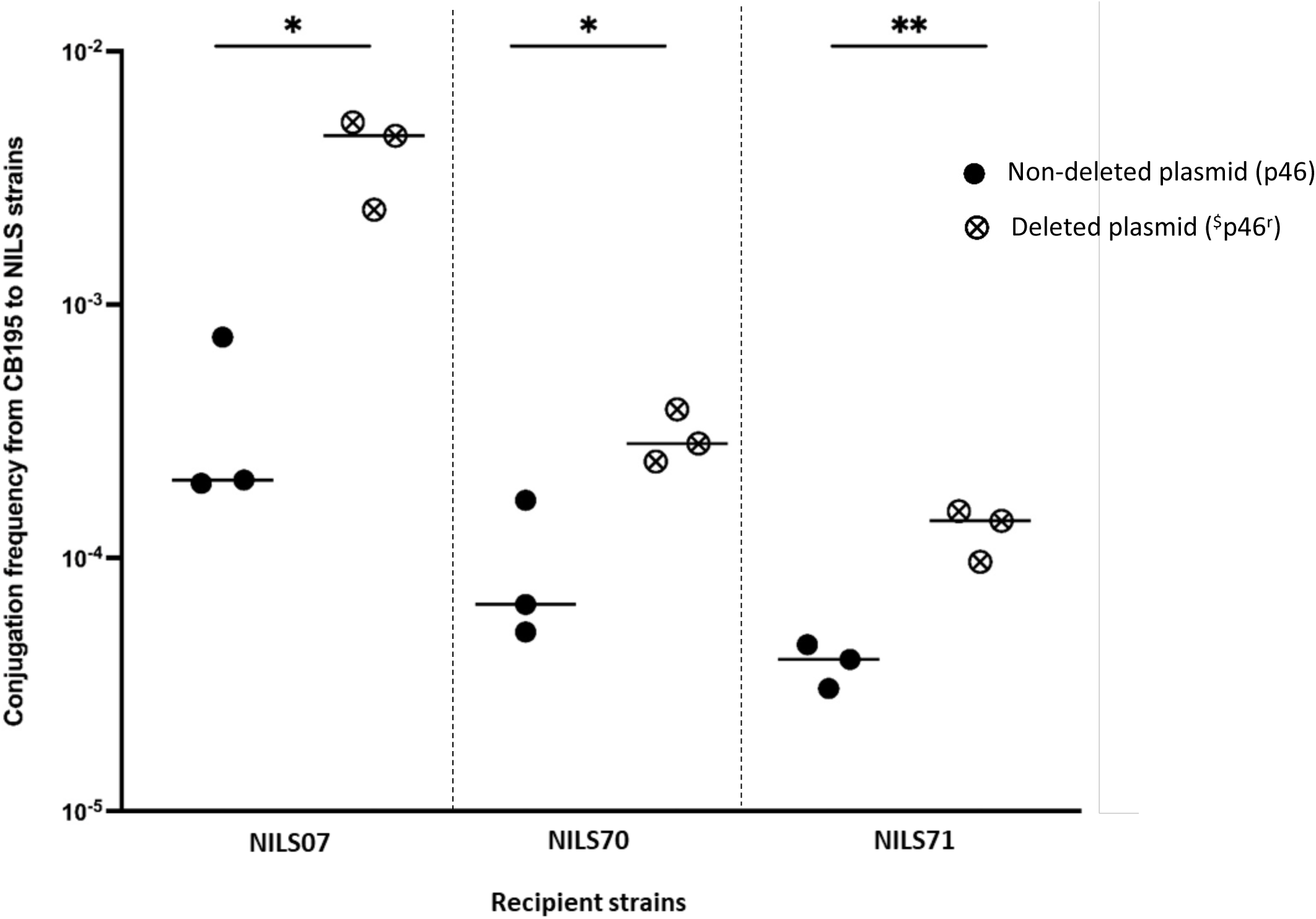
Scatter plot representation of the transfer frequency observed after conjugation of p46 (full circle) and p46^r^ (circle-cross), from ancestral donor strain CB195 to *E. coli* NILS strains. The dotted lines separate the different NILS strains. Each dot represents the result of one conjugation experiment. Each central line represents the mean of all the experiments. All experiments were performed in three biological replicates, with three technical replicates each. Asterisks indicate significant differences (*, *P* ≤ 0.05; **, *P* ≤ 0.01). ^$^ Here, p46^r^ plasmid was obtained from the ancestral donor strain CB195, and spontaneously exhibited rearrangements in the MDR-encoding region.

Independently of the background of the recipient strain, we observed a statistically significant increase in the transfer rate of p46^r^ plasmid compared to p46 plasmid.

### Effect of MDR-encoding region rearrangements on plasmid stability

To determine whether plasmid mutations or rearrangements could impact the maintenance of the p46 plasmid into new recipient hosts, we conducted an *in vitro* evolution experiment with six selected TC-NILS according to their plasmid rearrangements and chromosomal backgrounds: four NILS-p46^r^ presented large deletions in the MDR-encoding region (TC574, TC553, TC539 and TC544) (Figure 1) and two NILS-p46^nr^ showed no IncR backbone/aminoglycosides deletions (TC570 and TC549) (Figure 1 and Table S2). Three replicates of each TC-NILS were evolved in parallel in Lysogeny Broth (LB) with and without cefotaxime, by daily serial transfer. After 13 days (130 generations), we screened all evolved lineages for phenotypic resistance and for the presence of plasmid replicons (*repA* and *repB* genes, respectively).

In the presence of selective pressure (*i.e.* cefotaxime), no differences in phenotypic antimicrobial susceptibility and in replicon PCRs screening were detected for all evolved lineages. In contrast, we observed different evolutionary trajectories in antibiotic-free media, according to the background of the host strains. No differences (antimicrobial susceptibility and PCR-screening) were detected for TC-NILS07 and TC-NILS70, while cefotaxime-sensitive and *repA*^-^ colonies were identified in all three TC-NILS71, within one or three lineages (Table S5).

To further characterize all plasmid genome modifications, one clone of each evolved lineage, as well as cefotaxime-sensitive colonies from each TC-NILS71 lineage, were whole-genome sequenced. As expected, no plasmid genome modifications were detected in the evolved TC-NILS07 and TC-NILS70 clones. Loss of the entire p46 plasmid was observed in the evolved TC539 lineages, which was the TC-NILS71 harboring the most deleted ancestral plasmid. In addition, large deletions of the plasmid backbone genome, ranging from 80 kb to 108 kb and encompassing the *bla*_CTX-M-15_ gene as well as genes involved in the conjugative apparatus, were detected in evolved TC544 and TC549 lineages (Table S5).

### Growth-conjugation tradeoff of multidrug-resistant IncC plasmids according to MDR-encoding module rearrangements

The difference in growth dynamics of the clinical recipient strains prevented an accurate estimation of the way IncC plasmid rearrangements affected the vertical and horizontal transmission tradeoff (Figure S6). To overcome this limitation, we modeled plasmid dynamics, using the same *E. coli* K-12 MG1655 host strain and TC-NILS as new plasmid donor strains, enabling general comparison of plasmids excluding confounding host effects. We measured the MGRs of two cell populations (*i.e*. K-12 strains harboring p46^r^ and p46^nr^ plasmids), and determined the p46^r^ and p46^nr^ transfer rates from TC-NILS donor strains to the K-12 recipient strain.

With previously selected TC-NILS (four NILS-p46^r^, TC574, TC553, TC539 and TC544 and two NILS-p46^nr^, TC570 and TC549) (Figure 1 and Table S2), we conducted several independent transfers of each p46^nr^ and p46^r^ plasmid to obtain multiple independent TCs denoted as TC-K12 (referred as TC-K12_07_, TC-K12_70_ and TC-K12_71,_ where the number in subscript refers to the TC-NILS donor strains). After conjugation, we screened the absence of new conjugation-induced IncR backbone/aminoglycosides deletions in TC-K12 (for more details, see Table S6).

Second, for each pair TC-NILS donor/K-12 recipient, we measured the MGRs of five randomly selected TC-K12 harboring either p46^nr^ or p46^r^ plasmid (denoted as K12_x_-p46^nr^, K12_x_-p46^r-1^ or K12_x_-p46^r-2^, where the number x in subscript refers to the TC-NILS donor strains) immediately after conjugation. For further analysis, we pooled the MGRs of TC-K12, in respect of the donor strains chromosomal background and p46 plasmid deletion, and compared to the MGR of plasmid-free K-12 MG1655. We observed that plasmids imposed a growth burden on the bacterial cell compared to the plasmid-free K-12 MG1655 strain (coefficient *β*_1_ to *β*_3_ in Table S7, *P* < 0.0001). Except one (coefficient *β*_5_, *P* = 0.4886), the MGRs of TC-K12 strains harboring p46^r^ were either higher than those of TC-K12 strains harboring p46^nr^ (coefficient *β*_4_ = +0.026 div.h^-1^, *P* = 0.0126; *β*_6_ = +0.022 div.h^-1^, *P* = 0.038) (Table S7). Of note, the p46 deletion differed between the two p46^r^-carrying TC-K12_71_ which could explain the variable growth burden (coefficient *β*_5_ and *β*_6_) of the p46^r^ plasmids on the bacterial host (Figure 4A, for more details, see Table S7).

**FIG 4:**
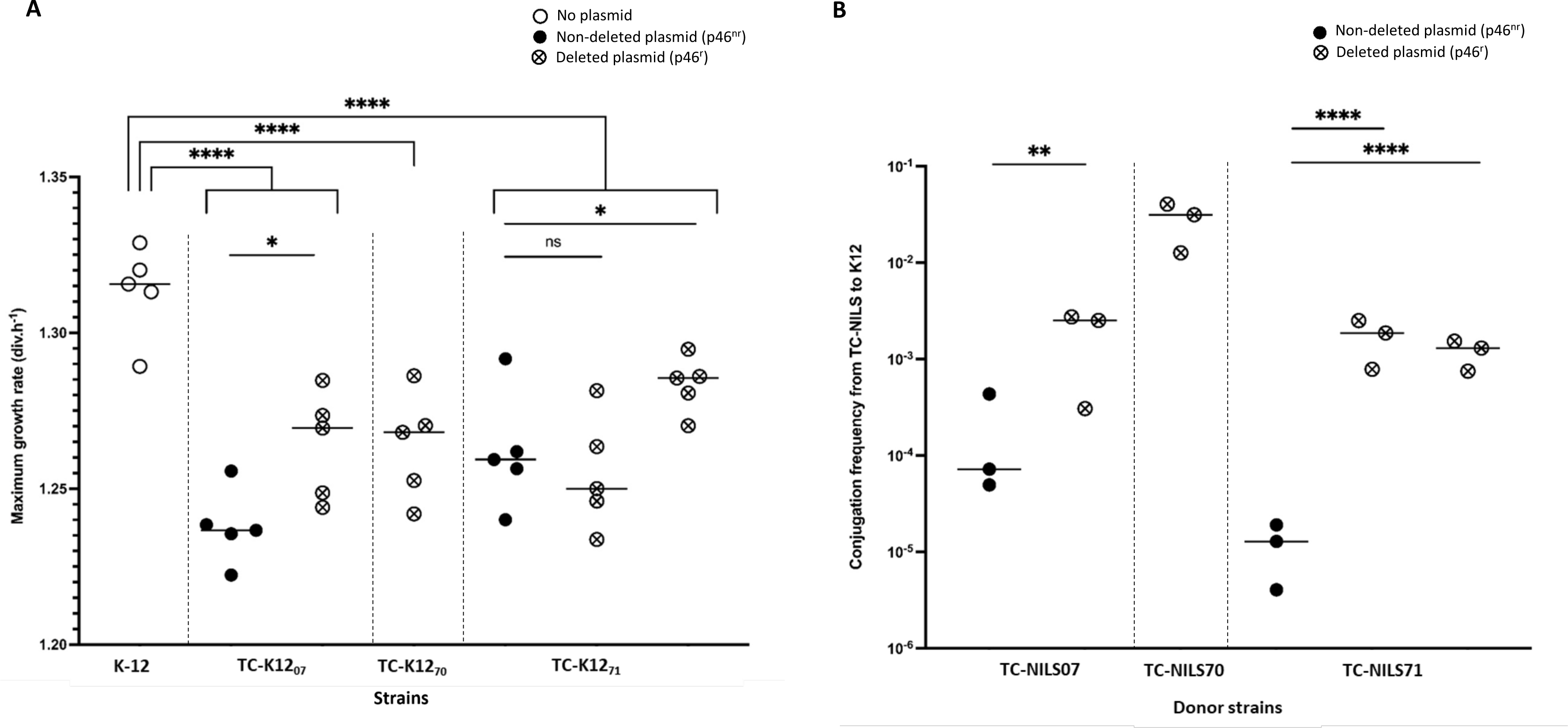
**(A)**, Scatter plot representation of the maximum growth rates (MGRs) of K-12 and TC-K12, obtained after conjugation of p46^nr^ (full circle) and p46^r^ (circle-cross) from TC-NILS to *E. coli* K-12 MG1655 srl::Tn*10* strain. The dotted lines separate the TC-K12 according to the chromosomal background of TC-NILS donor strains. Each dot represents the result of one experiment. Each central line represents the mean of all the experiments. All experiments were performed in three technical replicates. Asterisks indicate significant differences (*, *P* ≤ 0.05; ****, *P* ≤ 0.0001), ns = not significant. **(B),** Scatter plot representation of the conjugation frequency of p46^nr^ (full circle) and p46^r^ (circle-cross) from TC-NILS to *E. coli* K-12 MG1655 srl::Tn*10* strain. The dotted lines separate the TC-NILS donor strains according to their chromosomal background (*i.e.* NILS strains). Each dot represents the result of one conjugation experiment. Each central line represents the mean of all the experiments. All experiments were performed in three biological replicates, with three technical replicates each. Asterisks indicate significant differences (**, *P* ≤ 0.01; ****, *P* ≤ 0.0001), ns = not significant.

Third, we compared the transfer rate of p46^r^ and p46^nr^ from TC-NILS to K-12 MG1655 strain. We observed a significantly higher conjugation transfer frequency of the p46^r^ plasmid compared to the p46^nr^ plasmid. The conjugation frequency was 11 times higher from TC-NILS07 carrying p46^r^ than from TC-NILS07 carrying p46^nr^ (*P* = 0.0049) and respectively 155 and 115 times higher from TC-NILS71 carrying p46^r^ with two different deletions than from TC-NILS71 carrying p46^nr^ (*P* < 0.0001) (Figure 4B). We observed a significant difference in p46^r^ transfer rates according to the chromosomal background of TC-NILS donor strains (Kruskal-Wallis test, χ^df=2^ = 7.193, *P* = 0.0274). Overall, for all clinical strains from which they originated, the p46^r^ transfer rate was higher compared with p46^nr^ plasmids, regardless of their rearrangement, even though the conjugative capacity differed depending on the donor strain (Figure 4B).

Finally, we investigated the relationship between the conjugation frequency and the growth burden of plasmid carriage determined by MGRs measurements for the two cell populations (*i.e*. K-12 strains harboring p46^r^ and p46^nr^ plasmids), using the values and confidence intervals of MGR and conjugation frequency fitted by the linear models used for the analysis of these populations. Overall, we observed that the MDR-encoding module rearrangements increased the global fitness of the p46 plasmid, independently of chromosomal background of the donor strain. Indeed, the conjugative capacity of p46^r^ plasmids was systematically higher compared with p46^nr^ plasmids, without increased plasmid-induced growth burden (Figure 5).

**FIG 5:**
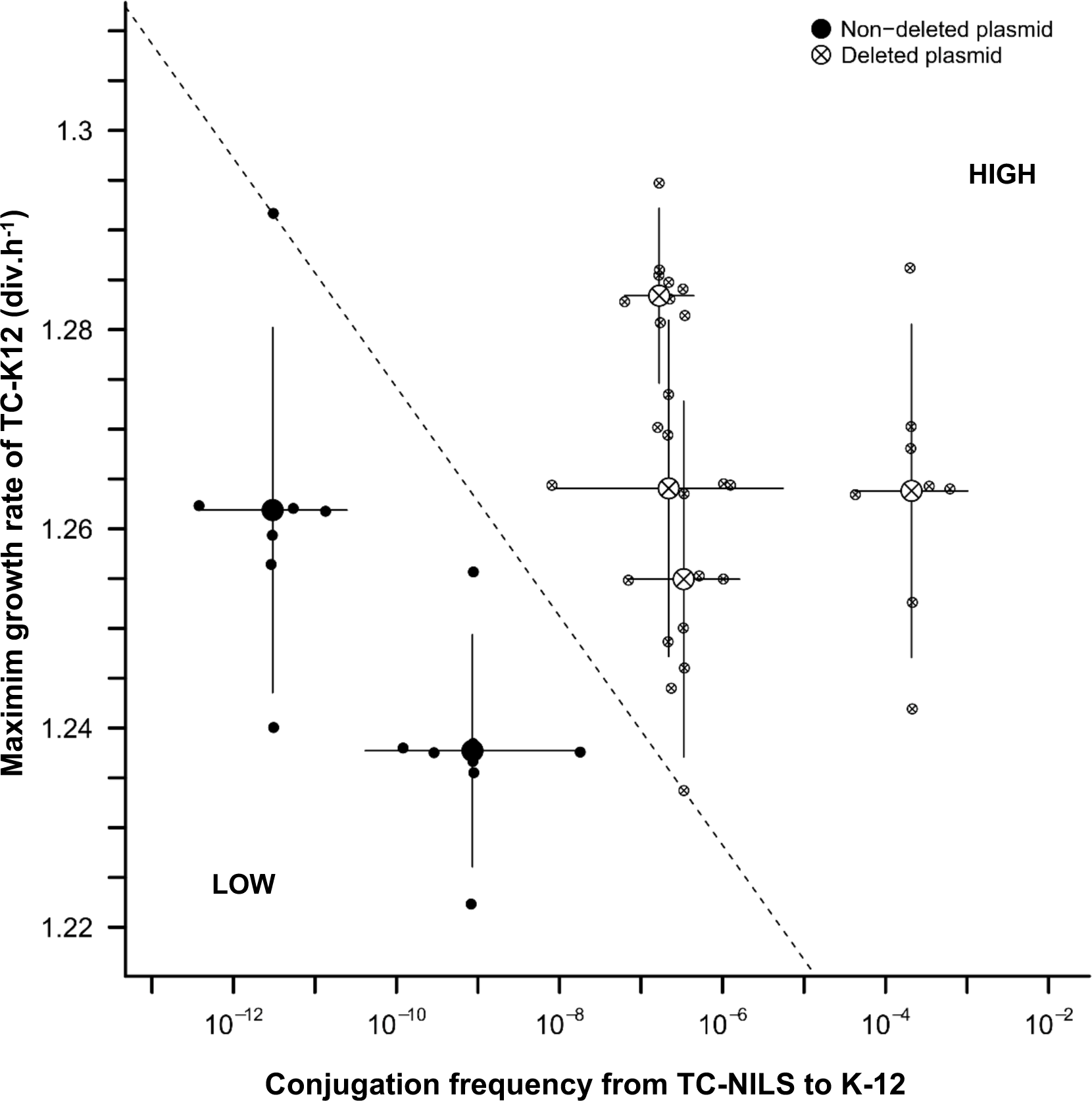
Two-dimensional representation of p46 plasmid dynamics. Relationship between the conjugation frequencies (x-axis) of p46^nr^ (full circle) or p46^r^ (circle-cross) from TC-NILS donor strains to K-12 MG1655 recipient strain, and maximum growth rates (MGRs) (y-axis) of the TC-K12 transconjugants obtained. All MGRs experiments were performed in five biological replicates, with three technical replicates, and all conjugation experiments were performed in three biological replicates, with three technical replicates each. The dotted line (which passes through the highest fitness value obtained in p46^nr^ plasmids and the lowest fitness value obtained in p46^r^ plasmids) separates their relative global fitness (low *vs* high).

## DISCUSSION

Understanding the evolutionary mechanisms underlying IncC plasmid success is a major challenge, due to its extremely broad host-range, its role in the emergence of resistance to medically important last-resort antimicrobials and its prevalence in environmental, animal and human bacteria (1, 2, 24). These plasmids share a closely related backbone and have diverged from a common ancestor through the acquisition of integrative elements encoding resistance genes (4, 8, 25). This study confirms our previous observations made in a laboratory adapted strain (15), and highlights the frequent genome rearrangements of a type 2 IncC plasmid immediately after its transfer in diverse clinical relevant strains. The MDR-encoding module rearrangements of this plasmid enhanced its transfer rate at the critical stage of entry into the host cell, without negative impact on its vertical transmission. Our results could explain the frequency of IncC plasmids among multidrug-resistant clinical bacteria, and their association with a wide variety of antimicrobial resistance (AMR) genes, including AMR genes derived from chromosomal beta-lactamases, such as *bla*_CMY_ (26), as these plasmids promote genomic diversity and their own genomic evolution.

The observed rearrangement rates of IncC plasmids ranged from none for p30 to high for p46, and concerned the MDR-encoding region in agreement with previously published observations of plasmid phylogeny (2, 3, 25). Both belonging to the IncC type 2 lineage, p30 and p46 plasmids differ in the content of their accessory regions and their integration sites in the plasmid core genome. All the genome rearrangements in p46, which contains more integrase genes than p30, consisted of IS*26*-mediated deletions. IS*26* uses two movement mechanisms, a classical replicative transposition and a very efficient and targeted conservative reaction, making any pre-existing copy of IS*26* an attractive target for additional IS*26*-assosiated segments (27, 28). This unique property explains the role of IS*26* in the creation of regions containing multiple copies of IS*26* interspersed with different DNA segments, such as complex resistance regions (29). IS*26* family is known as the most important contributor to the mobilization of AMR genes in Gram-negative bacteria (30–32), and actively contributes to the reorganization of plasmid genomes (33, 34).

In our model, p46 rearrangements mainly occurred immediately after conjugation and the conjugation process accelerated the mobilization of the transposons. Conjugative ssDNA transfer can trigger the SOS response and affect integrase gene expression in integrons, leading to rearrangement of gene cassettes (35). The expression of most IncC genes, including large operons involved in the conjugation process (36) and the bet/exo homologous recombination system, depends on AcaCD, a plasmid-borne master activator, also activating the expression of genes carried by distinct families of genomics islands (37–39). Previous studies have shown that conjugative entry of IncC plasmids induce the SOS response in recipient cells and served as helper conjugative plasmids to the mobilization and dissemination of genomics islands, via SOS induction, through the complex transcriptional regulatory network (37–39). Thus, the SOS response induction during early broad-host range IncC conjugative plasmid transfer might play a key role in its own genomic organization. Depending on the AcaCD binding sites present in the host cell genome, this could be an opportunity for extensive DNA exchange and acquisition of new resistance genes. In our experimental approach, we deliberately chose recipient strains without any antibiotic resistance genes, so that no acquisition of new AMR was observed in IncC plasmids during conjugation. The literature points to an environmental reservoir as the origin of the IncC in food-animal and human enteric bacteria (1), and to the rapid acquisition of resistance genes modules being associated with antibiotics used in animal agriculture and human therapy. Our observations highlight the ability of some IncC plasmids to undergo frequent genomic modifications immediately after the conjugation in a new host, which could explain their association with a wide range of critically important AMR genes.

Some of the p46 deletions observed in our experiments should have an impact on plasmid maintenance because they involve partition systems, important factors contributing to the stable inheritance of large, low-copy-number plasmids, such as IncC, in the bacterial population (40). However, p46 stability appeared variable after experimental evolution, affected exclusively in the absence of cefotaxime, and dependent on the background of the host strains. Non-deleted and deleted ancestral p46 plasmids were stably maintained in two of the three recipient strains without any additional modification. Hancock *et al.* have identified a crucial set of IncC backbone genes, adjacent to the *repA* gene, that are required for its stability (41) and probably contribute to p46 stability when the MDR-encoding region is deleted. After evolution, loss of the entire plasmid or backbone plasmid deletions, encompassing the *bla*_CTX-M_ cassette and a part of the conjugation apparatus, were only detected in evolved TC-NILS71 clones. These results are consistent with a long-term co-evolutionary adaptation between the plasmid and the recipient cells, as changes in conjugation rate and loss of nonessential AMR genes improve the cost of carriage for costly conjugative plasmids (16, 18, 42, 43). Taken together, these data suggest that rearrangements of the MDR-encoding module, associated with the conjugative entry of the IncC plasmid, represent an adaptive response different from the long-term compensatory evolution between plasmid and host.

In addition to the long-term fitness cost for their hosts, the initial energetic burden imposed by the newly acquired plasmid created a transient acquisition cost that manifested as a prolonged lag time, regardless of growth rate, especially for broad-host range plasmids (17). However, Ahmad *et al.* have proposed an evolutionary tradeoff whereby clones exhibiting longer lag times achieve faster recovery growth rates. Due to that lag/growth tradeoff, intermediate-cost plasmids may be evolutionarily favored compared to their lower and higher cost counterparts, in clonally heterogeneous populations (44). Genome modifications of IncC plasmid, immediately after conjugation in a new host, could have important ecological implications and could be important in understanding their prevalence in the environment.

Another important result in our study is the enhanced transfer rate of the IncC plasmid following MDR-encoding module rearrangements. Plasmid fitness is subject to an evolutionary tradeoff between horizontal and vertical transmission (16), and changing the conjugation rate is key to reducing the cost of plasmid carriage (18, 42, 45). However, one recent study showed a striking tolerance of several narrow-host range IncF plasmids to increased conjugation across multiple orders of magnitude, without significant growth burden (20). In our experiments, conjugation rates were higher for the deleted IncC plasmids, without any impact of the rearrangement on the fitness cost for recipient strains, during the period of plasmid establishment in a new bacterial host. This suggests that the systematic conflict between the two modes of plasmid transfer – vertical and horizontal transmission – is not immediately apparent after conjugation.

Several caveats apply to our results. First, we only used two types of type 2 IncC plasmids, which limits our understanding of the molecular mechanisms underlying their genomic changes and ecological success. However, the high structural stability of the genomic backbone of IncC plasmids suggests a direct role for MDR-encoding content in our observation. Recent progress has been made in understanding the molecular biology of broad-host range IncC plasmids, by describing a number of transcriptional regulators, critical to their broad-host dissemination (2, 4), and the high resilience to host defence systems upon entry in a new host by conjugation (46). Future investigations, using transcriptional analysis at the single-colony level, will help to fully understand the molecular mechanisms underlying genomic changes in the early stages following conjugative entry of the IncC plasmid. Second, we only tested a small number of clinical host bacteria. However, we chose natural isolates corresponding to phylogenetically distant isolates with different chromosomal backgrounds, and we obtained the same effect on p46 rearrangements and on the evolution of IncC plasmid fitness. Third, our observations were performed *in vitro* and under a single environmental condition. Some studies have described that, even in the absence of positive selection, the global fitness of the MDR conjugative plasmids could be improve by the presence of alternative hosts in multi-species bacterial communities (47, 48). Further investigations should be conducted in complex polymicrobial communities, such as microbiota, to determine at the population-level, the ecological impact of the IncC plasmid genome modifications on their persistence and dissemination.

In the present study, we propose an adaptive process, whereby vertical and horizontal transmission temporality of p46 improves, and which could favor the dissemination of the IncC plasmid in bacterial new hosts. Likewise, the capture of new antimicrobial genes by IncC plasmids could play a key role as a by-product of rapid adaption of incoming plasmid, in an environment involving co-resident plasmids interactions (*e.g.* complex microbial communities), or subject to antibiotic selection pressure. These findings highlight the importance of the IncC plasmid as a powerful tool for understanding the evolution of plasmid resistance.

## MATERIALS AND METHODS

### Bacterial strains and plasmids

We selected three *E. coli* clinical strains, previously whole-genome sequenced, for this study on the basis of their plasmid replicons and their sequence type (Table S1). They were selected among the NILS collection characterized by the absence of *rpoS* mutation (21) and were obtained during January 2013 from two clinical laboratories based in different teaching hospitals in Paris area, France. They were directly sampled from antibiograms on Mueller-Hinton (MH) plates, were sub-cultured only twice and were not subjected to stab storage prior to the extraction of genomic DNA and storage in glycerol at −80°C. NILS07 (accession number ERS13474783), strain of phylogroup A, ST10 according to Achtman multilocus sequence typing (MLST) scheme and serotype O105:H6 was isolated from feces. NILS70 (accession number ERS13474846), strain of phylogroup B2, ST12 (Achtman MLST scheme) and O4:H5 was isolated from urine. NILS71 (accession number ERS13474847), strain of phylogroup B2, ST1154 (Achtman MLST scheme, single locus variant of ST73) and O2:H1 was isolated from urine. None of these strains carried antimicrobial resistance gene based on whole-genome sequence analysis. To generate spontaneous rifampicin-resistant (Rif^R^) mutants, wild-type NILS strains were incubated in LB + 1x the Minimum Inhibitory Concentration of rifampicin overnight at 37°C, and spread (300 µL/plate) on LB agar medium containing rifampicin (30 mg/L). The plates were then incubated at 37°C for 24 hours, so that the background remains as close as possible to the original strain. In the end, NILS07 showed a mutation at codon 526 (CAC to C<u>T</u>C), and NILS70 and NILS71 at codon 513 (CAG to C<u>T</u>G) of the *rpoB* gene conferring resistance to rifampicin.

We selected two broad-host range ESBL-encoding IncC plasmids used in previous studies, pRCS30 (GenBank accession number LT985224) and pRCSC46 (PRJEB70418-ID17151429), which we will referred to as p30 and p46 respectively, for simplicity (15, 22) (Table S1 and Figure S2). p30 (157,836 bp) carries two resistance islands, inserted at two different integration spots: on the one hand, a *bla*_CTX-M-14_ transposition unit and, on the other hand, a group of four resistance gene (*sul2, strA-strB, ermMI*, and *bla*_TEM-1_), named the ARI-B island by Harmer and Hall (2). This plasmid originated from an *E. coli* strain of phylogroup F and ST721 (MLST Pasteur scheme) (strain CB513) isolated from a case of bacteriemia in an adult patient in 1999 (Paris area, France). p46 (246,268 bp) carries a *bla*_CTX-M-15_ transposition unit which is inserted downstream of a methylase encoding gene. On this plasmid, a large region of 122,219 bp is inserted into the *rhs* gene. This region contains two resistance modules (an IS*26*-mediated *aacC2-aph-sul2* cassette followed by a *dfrA-sul1* cassette), a partial IncQ1 replicase and the complete backbone of an IncR plasmid (17,000 bp). The plasmid is thus a multireplicon composite plasmid, IncC-IncR, which contains two complete replicons. This plasmid originated from an *E. coli* strain of phylogroup A and ST718 (MLST Pasteur scheme) (strain CB195) isolated from a case of fecal carriage in an adult patient in 1999 (Paris area, France).

A third plasmid, derived from p46 (denoted as p46^r^), was obtained in a fraction of the bacterial population of the ancestral strain CB195, after an *in vitro* experimental evolution (80 to 130 generations), and had spontaneously lost the IncR backbone and its flanking regions, as well as the aminoglycosides resistance module (*aacC2*-*aph-sul2* cassette).

### Mutation rate assays

We estimated mutation frequencies of the isolates by monitoring their capacity to generate mutations conferring resistance to rifampicin or fosfomycin (49). We used *E coli* strain M13, a strong mutator because of a defective mismatch repair system due to MutS inactivation (*mutS* mutant) as a positive control (49), and a wild-type *E. coli* strain K-12 MG1655, as a negative control.

From a glycerol stock, the strains were plated on LB agar and grown overnight at 37°C. We used randomly selected colonies for independent mutation rate assays. They were grown overnight in 10 mL of LB at 37°C under constant stirring at 200 rpm. Next, we transferred 10 µL of the overnight cultures into 990 µL of fresh LB and 100 µL of these cultures were transferred in 10 mL of fresh LB and incubated overnight in the same conditions. Then, we plated appropriate dilutions of the overnight cultures on LB agar without antibiotics, LB agar containing rifampicin (250 mg/L) and LB agar containing fosfomycin (50 mg/L). We counted the rifampin-resistant mutants and fosfomycin-resistant mutants after 24 h at 37°C. We calculated the mutation frequency as the number of mutants obtained on LB with antibiotics divided by the total number of bacteria on LB without antibiotics. We performed the mutation rate experiments in three biological replicates with at least two technical replicates each.

### Conjugation assays

First, we performed independent transfer assays by conjugation of the three ESBL-encoding plasmids (p30, p46, and p46^r^) from the donors (CB513 and CB195, respectively) to *E. coli* NILS strains (NILS07, NILS70, and NILS71), made resistant to 250 mg/L rifampicin, as the recipient strains. From a glycerol stock, the donor and recipient strains were plated on LB agar and grown overnight at 37°C. We used randomly selected colonies of donor and recipient strains for independent conjugation assays. Donors and recipients were grown overnight in 10 mL of LB at 37°C under constant stirring at 200 rpm. Next, 10 µL of the culture was transferred into 10 mL of fresh LB and incubated for 3 h under the same conditions to obtain an exponential-phase population (OD_600_ = 0.4-0.5). For mating experiments, we mixed 1 mL of culture of the donor strain and 1 mL of culture of the recipient strain (1:1 ratio) and we immediately plated 1.8 mL (9 spots of 200 µL) of mating culture on LB agar without antibiotic at 30°C. After 3 hours of incubation, we resuspended cultures in 5 mL of fresh LB. Then, resuspended mating cultures were plated on LB agar containing rifampicin (250 mg/L) and cefotaxime (4 mg/L) and incubated overnight at 37°C for TCs selection (referred as TC-NILS).

Second, we performed independent transfer assays by conjugation of ESBL-encoding plasmids to *E. coli* recipient strain K-12 MG1655 srl::Tn*10,* resistant to tetracycline. For this second conjugation, we selected as donor strains six TC-NILS obtained from our previous conjugation experiments, according to plasmid rearrangements and chromosomal background strain. We used the same protocol as previously, except that we selected the new TCs (referred as TC-K12) using LB agar containing tetracycline (20 mg/L) and cefotaxime (4 mg/L) (Figure S7).

We calculated the conjugation frequency as the number of TCs divided by the total number of bacteria. We performed the conjugation experiments in three biological replicates with three technical replicates each.

### Antimicrobial susceptibility testing

We performed antimicrobial susceptibility testing using the disk diffusion method according to the European Committee on Antimicrobial Susceptibility Testing (EUCAST) guidelines (http://www.eucast.org). The list of antimicrobial compounds that were tested included: ampicillin, amoxicillin-clavulanic acid, ticarcillin, ticarcillin-clavulanic acid, piperacillin, piperacillin-tazobactam, temocillin, cefalexin, cefoxitin, cefixime, ceftazidime, cefotaxime, cefepim, aztreonam, ertapenem, meropenem, imipenem, amikacin, gentamicin, tobramycin, netilmicin, streptomycin, trimethoprim, cotrimoxazole, nalidixic acid, ofloxacin, ciprofloxacin, fosfomycin, tetracycline, rifampicin, chloramphenicol.

### Plasmid rearrangement screening

To determine the rearrangement frequencies of the p46 plasmid after conjugation, we screened TCs for phenotypic aminoglycosides resistance (see above for the antimicrobial susceptibility testing), and for the presence of the gene encoding IncR replicase (*repB* gene). We tested TC-NILS obtained from the first conjugation assay between CB195 and NILS strains and TC-K12 obtained from the second conjugation assay between TC-NILS and K-12 MG1655 strain. We obtained the total DNA by boiling lysis method exposing cells to 100°C for 10 min and clarifying the lysate preparations by centrifugation at 15,000 x g for 10 min. We determined the *repB* status on DNA extracts by a specific PCR with a set of primers for IncR-F (5’ TCG CTT CAT TCC TGC TTC AGC 3’) and IncR-R (5’ GTG TGC TGT GGT TAT GCC TCA 3’). A 251 bp fragment internal to IncR was amplified. We analyzed the amplification products by 1.6 % agarose gel electrophoresis. We also used the PCR-Based Replicon Typing (PBRT) 2.0 Kit (Diatheva, Fano, Italy) to determine replicon content of some TCs, according to the manufacturer’s recommendations (50–52).

### Plasmid stability

The six TC-NILS, selected according to their plasmid rearrangements and chromosomal background strains, were inoculated into 5 mL of fresh LB and grown under constant shaking at 200 rpm. After overnight incubation at 37°C, 5 µL of the culture was inoculated into three tubes containing 5 mL of fresh LB and three additional tubes containing 5 mL of fresh LB plus cefotaxime (2 mg/L). At this step, the 36 cultures were considered the ancestors of each lineage. Next, the 36 cultures were propagated by daily serial transfer, with a 1:1000 dilution (5 µL of culture grown overnight diluted in 5 mL of fresh LB with and without cefotaxime), for 13 days (corresponding to 10 generations per days; 130 generations in total) and incubated overnight at 37°C under constant stirring at 200 rpm. Every 3 or 4 days of the evolution assay, a culture sample from each lineage was collected and stored at -80°C. At day 13, whole population of each lineage was tested for phenotypic antimicrobial susceptibility and two specific PCRs targeting *repB* and *repA* genes, encoding for IncR and IncC replicase genes respectively, were performed. We determined the *repA* status on DNA extracts by a specific PCR amplifying a 465-bp fragment using the following primers: IncC-F (5’-GAGAACCAAAGACAAAGACCTGGA-3’) and IncC-R (5’-ACGACAAACCTGAATTGCCTCCTT-3)’. The *repB* status was determined as in the plasmid rearrangement screening.

### Whole-genome sequencing

We selected TC-NILS from conjugation of p30 and p46 plasmids and evolved TC-NILS for whole-genome sequencing (WGS) using Illumina technology. Briefly, DNA was prepared for indexed, paired-end sequencing on the Illumina MiSeq system (Integragen, MA, USA). DNA samples were extracted using the EZ1 DNA Tissue Kit (Qiagen, Courtaboeuf, France). The DNA was quantified using the Qubit Fluorometer (ThermoFisher Scientific, Asnières sur Seine, France). DNA libraries were prepared and indexed, and tagmentation was processed using the Nextera DNA Flex library Prep kit (Illumina, USA). The sample concentrations were normalized in equimolar concentration at 1 nM. The pooled paired-end libraries were sequenced to a read length of 2 by 300 bp with Illumina MiSeq reagent kit. The genomes were sequenced at an average depth of 50X.

### DNA sequence analysis

Illumina genome sequence reads were assessed for quality using FastQC v0.11.8 and subsequently trimmed with a cutoff phred score of 30 using TrimGalore v0.4.5. All genomes were *de novo* assembled using SPAdes v.3.13.1 software and analysed with an in-house bioinformatics pipeline (https://github.com/iame-researchCenter/Petanc) adapted from Bourrel *et al*. (53). Briefly, genome typing was performed including MLST determination, phylogrouping, and serotyping. The pipeline was also used to get information about plasmid replicons, resistance and virulence genes, the prediction of a plasmid sequence or a chromosome sequence for each contig using PlaScope (54). For each sample, sequenced reads were mapped to a reference sequence to analyze the mutations and rearrangements. For reference chromosomal sequences, we used the NILS sequences extracted *de novo*. For reference plasmid sequences, we used the plasmid sequences annotated on the MicroScope plateform (55). We identified point mutations and rearrangements using the open computational pipeline Breseq v0.27.1 (23). All SNPs identified in the ancestor strains that were sequenced were subtracted from the sequences of the TCs.

### Individual bacterial fitness assays

We determined bacterial fitness, defined here as the Maximum Growth Rate (MGR), from growth experiments by OD measurements up to 24 hours of growth. From a glycerol stock, the strains were plated on LB agar and grown overnight at 37°C. We used randomly selected colonies for independent growth assays. They were grown overnight in 10 mL of LB at 37°C under constant stirring at 200 rpm. Next, 200 µL of a 10000-fold dilution of each overnight cultures were incubated in a 96-well flat-bottom microtiter plate (Corning, ME, USA) at 37 °C for 24 hours and were subject to shaking during 60 seconds every two minutes. Optical density (DO 600 nm) was measured automatically every 300 seconds with a microplate reader (Infinite F200-Pro®, Tecan). The measurements were collected using a Python script developed in our laboratory. Each growth curve was characterized by three parameters: the lag time in minute (the time taken to reach the MGR), the MGR in sec^−1^ and the biomass corresponding to the OD_600_ in stationary phase after 24 h of growth. We performed the fitness assays in four biological replicates with three technical replicates each.

### Statistical analyses

For the first conjugation assay in NILS strains, the statistical tests were performed with GraphPad Prism version 9.5.1 software. Distribution normality was tested with a Shapiro-Wilk test and either two-tailed unpaired Student t tests or a Mann-Whitney tests were then performed depending on normality. To compare the rearrangements rates, a chi-squared test was used. Differences in MGRs (of TC-NILS and TC-K12) and conjugation frequency in K-12 MG1655 were analyzed using the R software. Linear models were used, with a response variable *Y(x)* (either the MGR of the bacteria or the conjugation frequency of the plasmid from the TC-NILS) computed as a linear combination of coefficients:

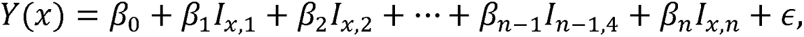

with □ the error, *β*_0_ the intercept, *β*_i_ the *i*^th^ coefficient of the model and *a*_x,i_ the element of the contrast matrix indicating how coefficient *β*_i_ affected the value of *Y(x)* (Table S4 and S7).

The contrast matrix characterizing the MGR of TC-NILS was defined such that specific coefficients tested for the difference between given bacteria: *β*_1_ and *β*_2_ characterized differences between TC-NILS07 and respectively TC-NILS70 and TC-NILS71, *β*_3,_ *β*_4_ and *β*_5_ characterized differences between bacteria with or without plasmid, respectively among TC-NILS07, TC-NILS70 and TC-NILS71, *β_6_*, *β*_7_ and *β*_8_ characterized differences between p30 and p46^nr^, respectively among TC-NILS07, TC-NILS70 and TC-NILS71, *β*_9,_ *β*_10_ and *β*_11_ characterized differences between p46^nr^ and p46^r^, respectively among TC-NILS07, TC-NILS70 and TC-NILS71.

The contrast matrix characterizing the MGR of TC-K12 was defined such that *β*_1_, *β*_2_ and *β*_3_ characterized the differences between plasmid-free K-12 and respectively TC-K12_07_, TC-K12_70_, TC-K12_71_, *β_4_* the difference between K12_07_-p46^nr^ and K12_07_-p46^r-1^, *β*_5_ and *β*_6_ the difference between K12_71_-p46^nr^ and respectively K12_71_-p46^r-1^ and K12_71_-p46^r-2^.

The contrast matrix characterizing the conjugation frequency was defined such that *β*_1_ and *β*_2_ characterized differences between transfers to TC-K12_07_ and respectively those to TC-K12_70_ and to TC-K12_71_, *β*_3_ characterized the difference between transfers to K12_07_-p46^nr^ and to K12_71□_p46^r□1^, *β_5_* and *β*_6_ the difference between transfers to K12_71_-p46^nr^ and respectively to K12_71_-p46^r□1^ and to K12_71_-p46^r□2^. While values of *β*_1_ in the models of MGR correspond to the actual differences in div.h^-1^, the values of *β*_1_ in this model correspond to the log-ratio between the conjugation frequencies.

For all analyses, *P* values of < 0.05 were considered significant.

## ACKNOWLEDGMENTS

We thank Marie-Florence Bredeche for the K-12 MG1655 srl::Tn*10* strain. We also thank Olivier Clermont, Linda Houhamdi and Marie Petitjean for technical assistance. We are grateful to Olivier Tenaillon for helpful discussion.

This work was partially supported by a grant from the Fondation pour la Recherche Médicale to Erick Denamur (grant number DEQ20161136698). The funders had no role in study design, data collection, and interpretation, or the decision to submit the work for publication.

## DATA AVAILABILITY

All reads have been deposited at the European Nucleotide Archive (project accession number PRJEB70418 and PRJEB56162, ID samples from ERS17121429 to ERS17121457, and ERS13474783, ERS13474846 and ERS13474847.

