## Supplementary material for "IncC plasmid genome rearrangements influence the vertical and horizontal transmission tradeoff in *Escherichia coli*": Fig S1 - S7

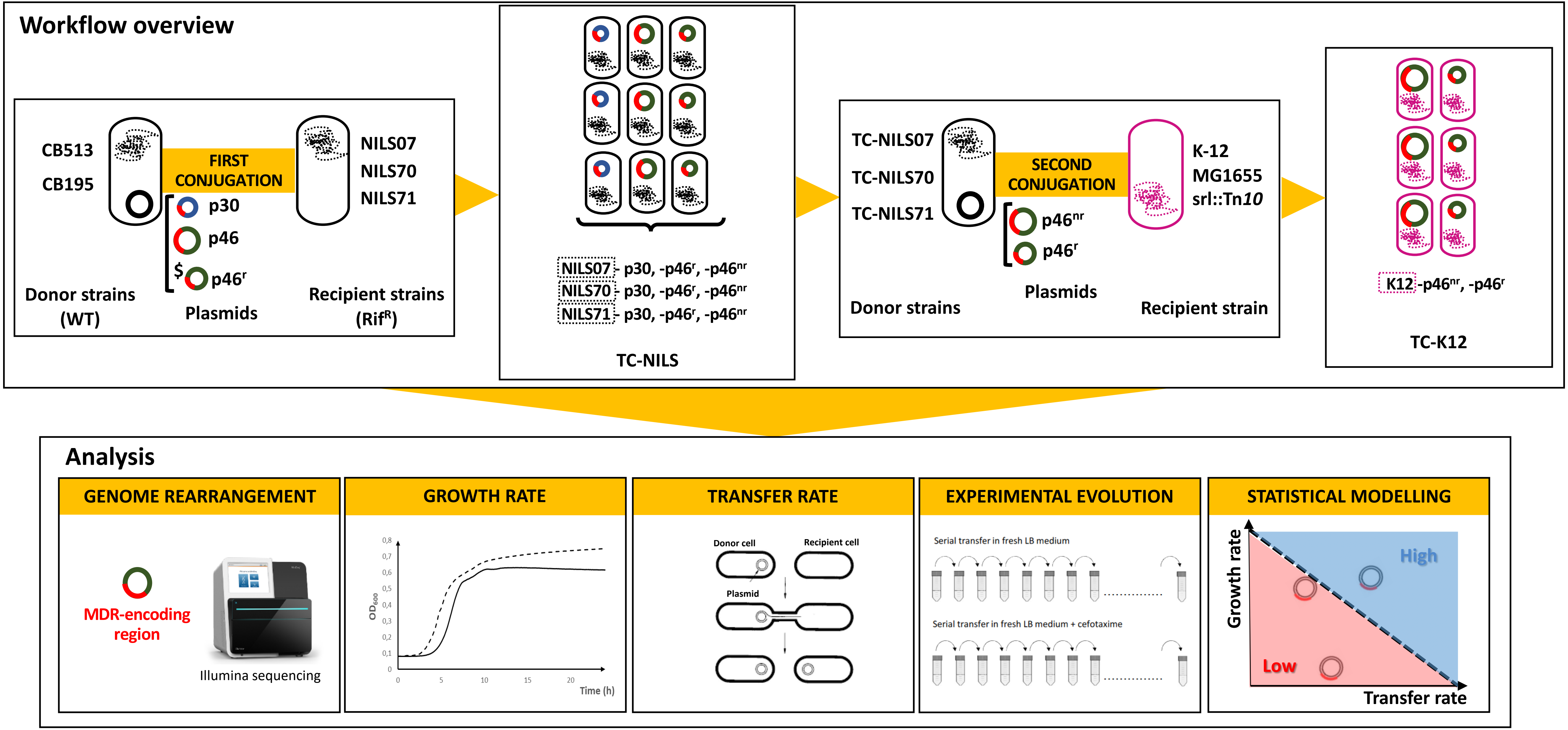


**FIG S1:** Workflow overview. First, multiple independent transfers of two plasmids (p30 and p46 from CB513 and CB195 wild type (WT) donor strains, respectively) in 3 clinical *E. coli* recipient strains (NILS07, NILS70, and NILS71; made resistant to rifampicin ($Rif^R$)) were performed to obtain multiple NILS transconjugants (denoted as TC-NILS). $^{\$}$, Independent transfers of p46 plasmid, which spontaneously exhibited rearrangements into the MDR-encoding region and selected from the ancestral donor strain CB195 (designated $p46^r$), were also performed in the 3 *E. coli* NILS recipient strains. Rearrangements that occurred during conjugation were analyzed by whole genome sequencing and their frequency was evaluated. Growth rates of the transconjugants were measured and the transfer rates of the various plasmids were determined. An *in vitro* experimental evolution in LB, without and with cefotaxime, was performed to evaluate the maintenance of the $p46^r$ plasmid in TC-NILS. Second, conjugation between previously formed TC-NILS (harboring $p46^r$ or non-deleted p46 plasmid, designated $p46^{nr}$) and K-12 MG1655 were performed to obtain multiple K-12 transconjugants (denoted as TC-K12). Finally, statistical modelling was used to determine the impact of rearrangements on vertical and horizontal plasmid transmission tradeoff.

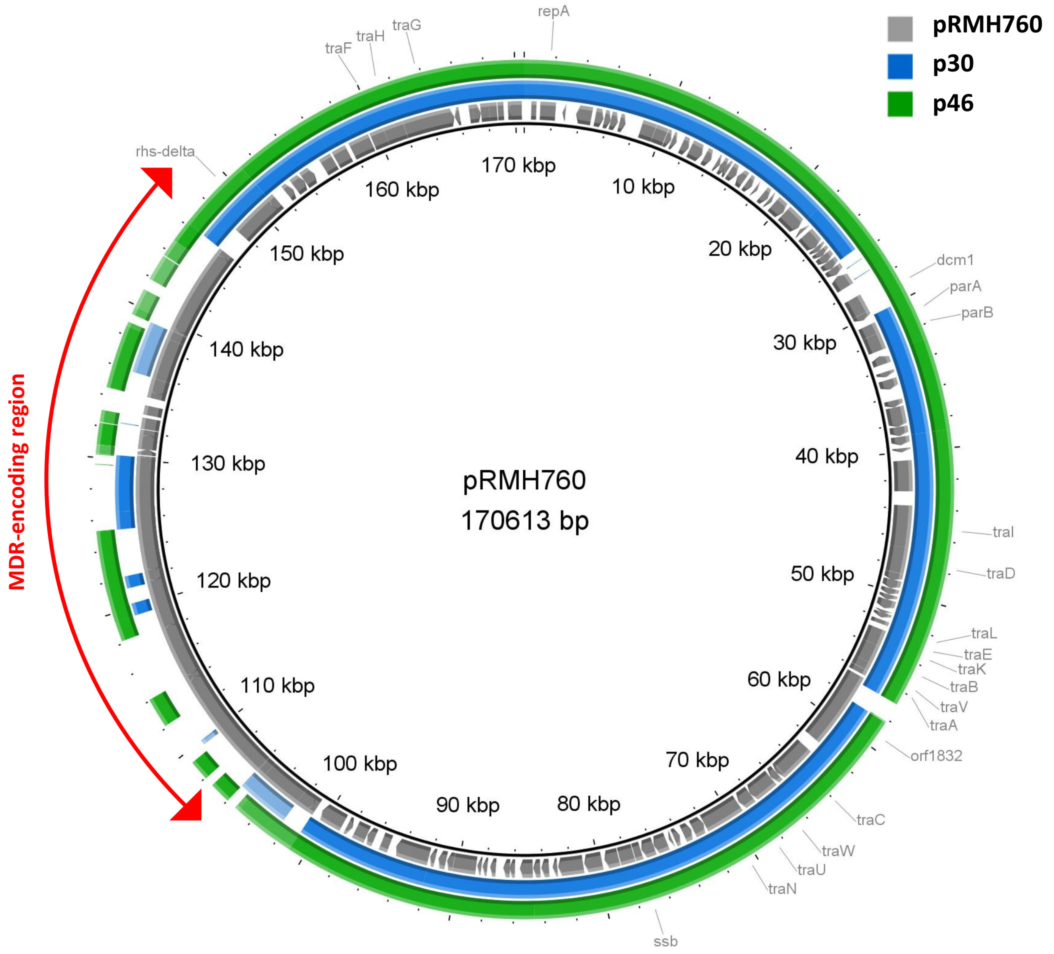


**FIG S2:** Comparative map of p30 (157,836 bp, type 2 IncC plasmid) and p46 (246,268 bp, type 2 IncC-IncR multireplicon composite plasmid) plasmids, with plasmid pRMH760 (170,613 bp, type 1a IncC plasmid, GenBank accession number KF976462) used as reference, generated using Blast Ring Image Generator (BRIG) diagram. The outer circle (green) represent the genome of the p46 plasmid, the central circle (blue) represent the genome of the p30 plasmid and the inner circle (gray) represent the genome of the reference plasmid pRMH760. Open reading frames are shown as arrow indicating the direction of transcription. The MDR-encoding region is indicated by double arrow.

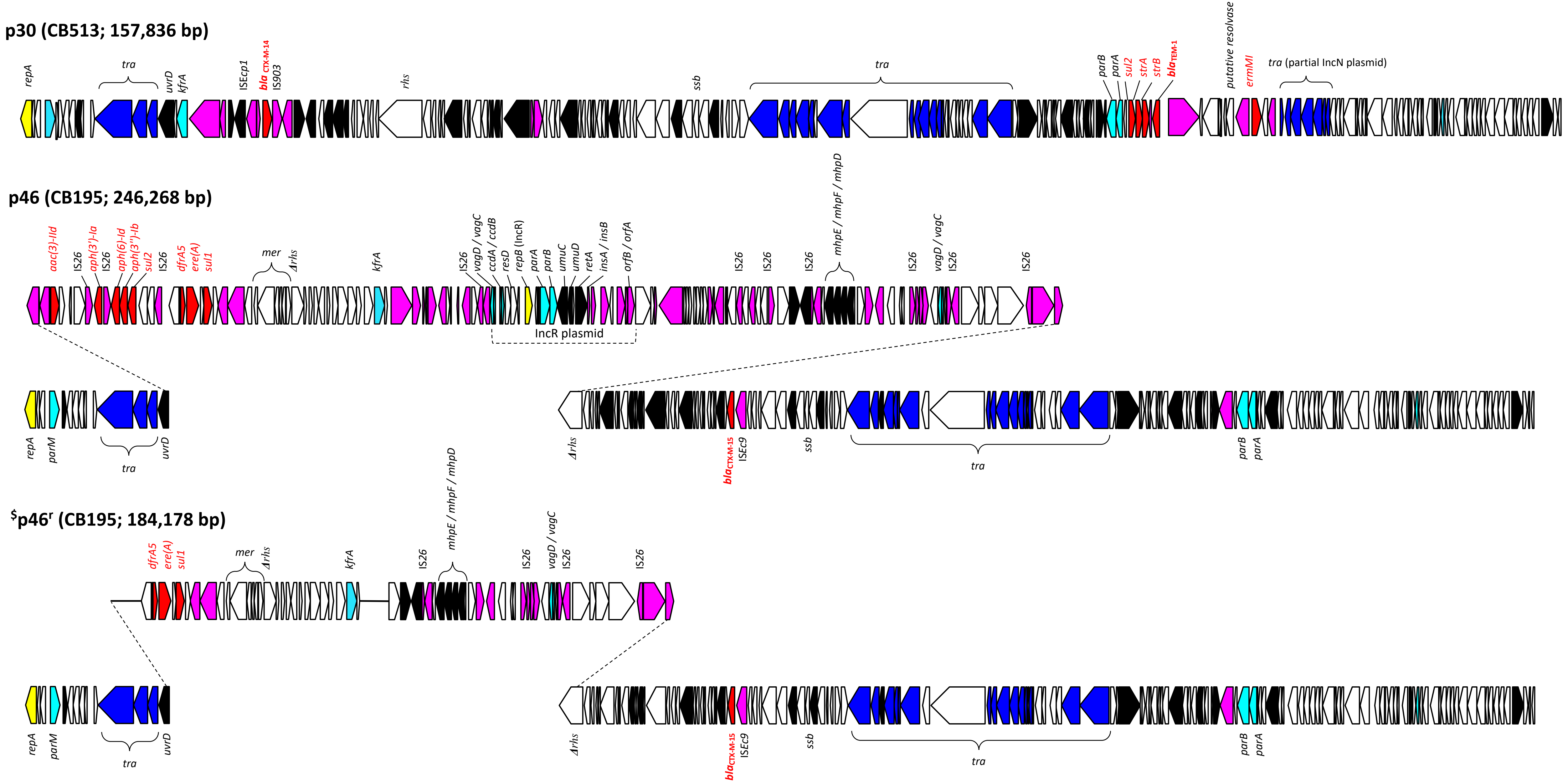


**FIG S3:** Linear maps of p30 (IncC, from CB513) and p46 (multireplicon composite plasmid IncC-IncR, from CB195) ancestral plasmids. Open reading frames are shown as arrows indicating the direction of transcription. Dark blue, plasmid transfer; yellow, replication; light blue, plasmid maintenance; red, resistance; black, metabolism; pink, mobile elements and white, hypothetical proteins. $ Rearranged p46 (denoted as p46r) plasmid (IncC), from CB195 ancestral donor strain and which spontaneously exhibited rearrangements into the MDR-encoding region, is also represented. Deleted regions in p46r compared to p46 (IncR backbone genes and their flanking regions, and the *aacC2-aph-sul2* cassette which confers resistance to aminoglycosides) are represented by a solid black line.

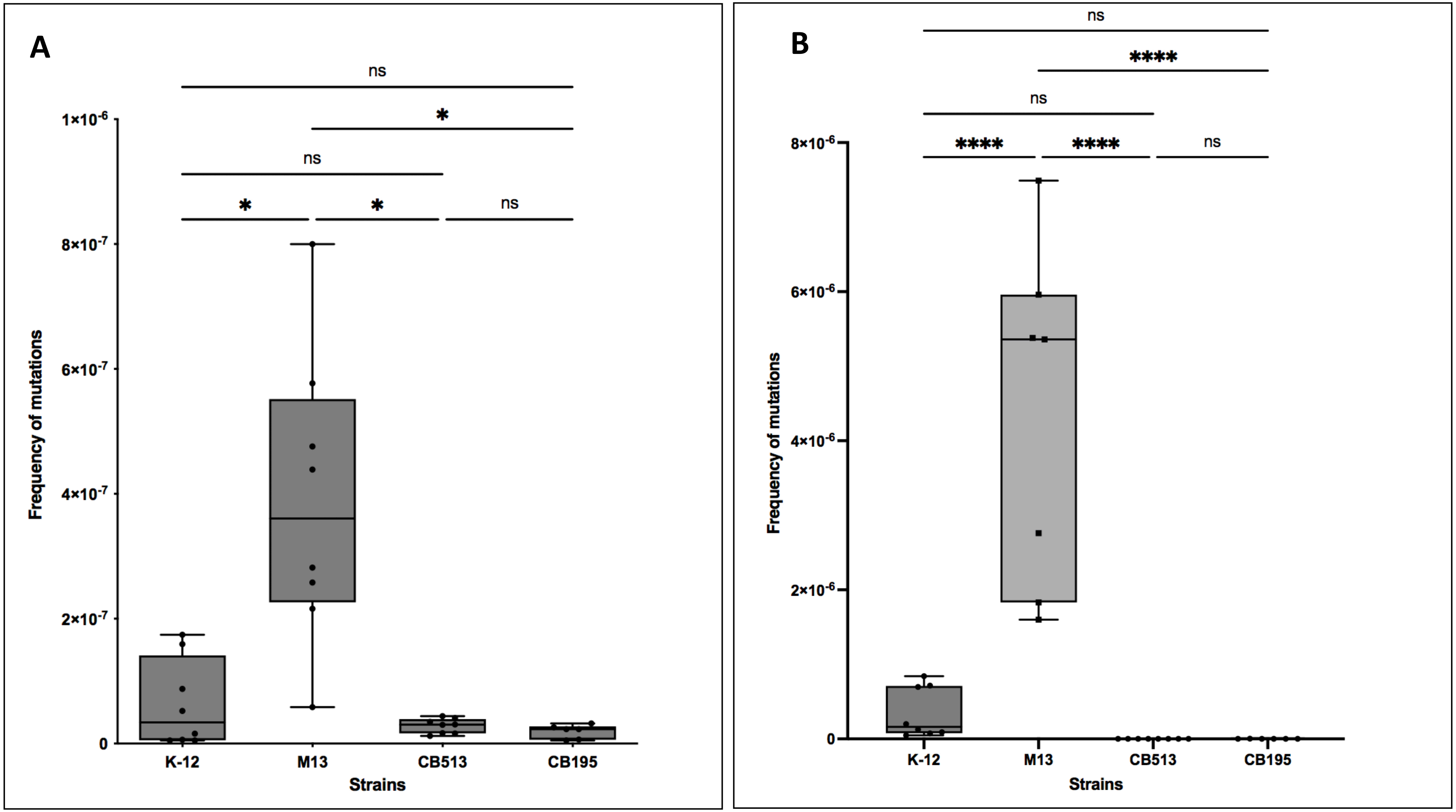


**FIG S4:** Boxplot representation of mutation rates of each *E. coli* donor strains (CB513 harboring p30 and CB195 harboring p46). (A) Frequency of rifampicin resistance acquisition. (B) Frequency of fosfomycin resistance acquisition. For each box, the central mark indicates the median, and the bottom and top edges of the box indicate the 25th and 75th percentiles, respectively. Each point corresponds to one measurement per clone. *E. coli* M13 and K-12 MG1655 are mutator (*mutS*-deleted) and non mutator controls, respectively. Asterisks indicate significant differences (*, $P \leq 0.05$; ****, $P \leq 0.0001$), ns = not significant.

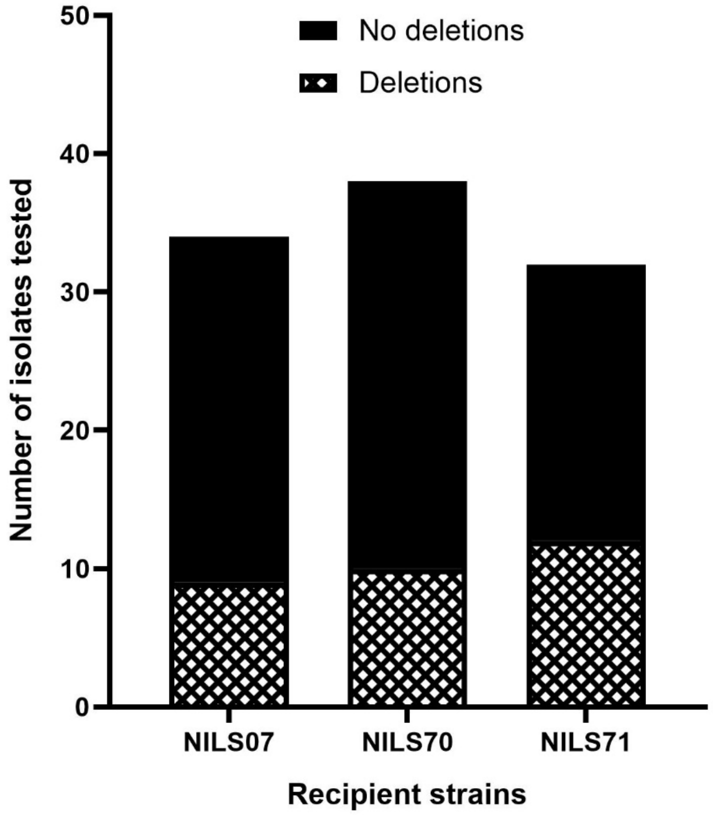


**FIG S5:** Share of isolates presenting plasmid IncR backbone/aminoglycosides deletions after conjugation of the donor strain CB195 in 3 *E. coli* clinical strains (NILS07, NILS70, and NILS71)

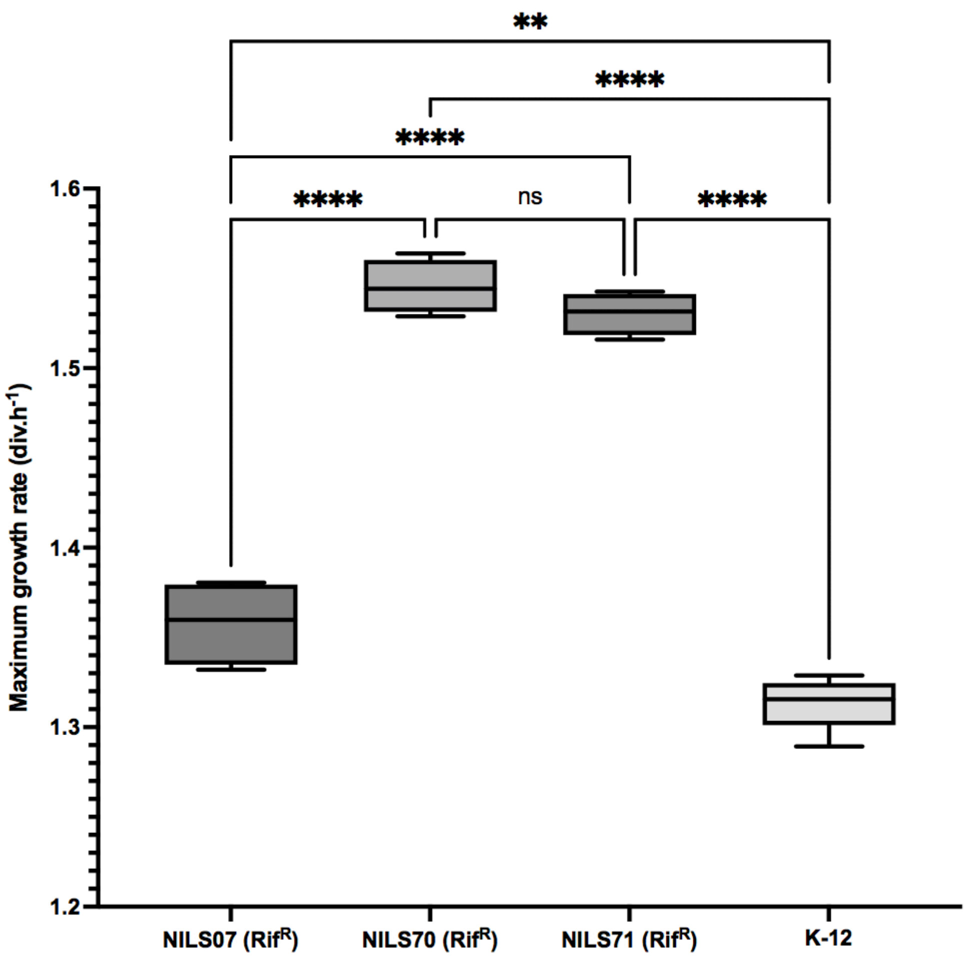


**FIG S6:** Box plot representation of the maximum growth rates (MGRs) of each recipient strains used in the study : NILS07, NILS70, NILS71, and K-12 MG1655. For each box, the central mark indicates the median, and the bottom and top edges of the box indicate the 25th and 75th percentiles, respectively. Asterisks indicate significant differences (**, $P \leq 0.01$; ****, $P \leq 0.0001$), ns = not significant.

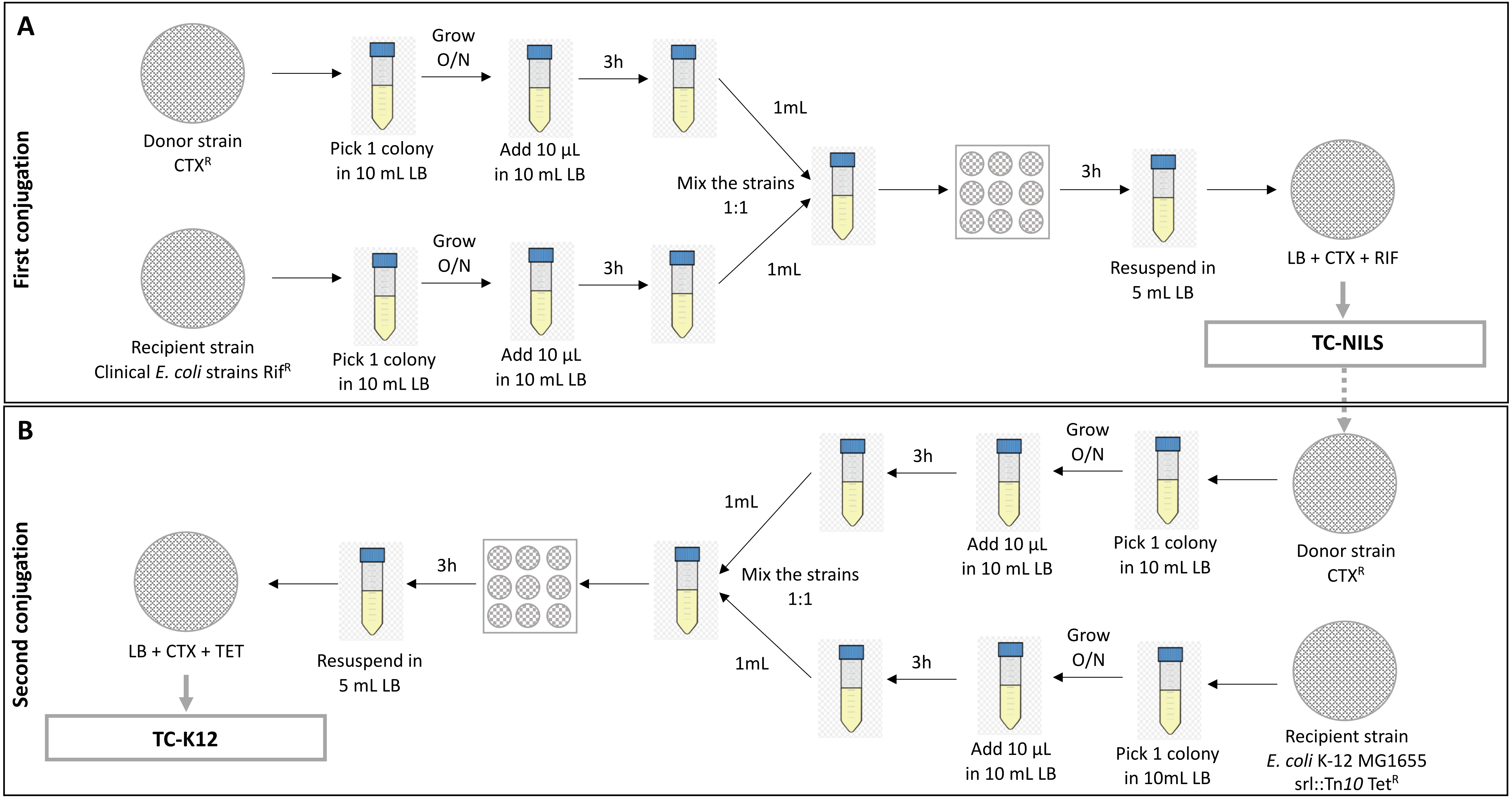


**FIG S7:** Design of the experimental conjugation assay. (A) First conjugation of 3 ESBL-encoding plasmids (p30 from CB513 donor strain, and p46 or p46$^{r}$ from ancestral CB195 donor strain) in 3 *E. coli* clinical strains (NILS07, NILS70, and NILS71) resistant to rifampicin (Rif$^{R}$). (B) Second conjugation of 6 TC-NILS (harboring p46$^{r}$ or p46$^{nr}$ plasmids, with or without IncR backbone/aminoglycosides deletion) obtained from conjugation A in K-12 MG1655 srl::Tn*10* strain, resistant to tetracycline (Tet$^{R}$). CTX = cefotaxime, RIF = rifampicin, TET = tetracycline.
